# A new spatial multi-omics approach to deeply characterize human cancer tissue using a single tissue section

**DOI:** 10.1101/2024.10.03.616510

**Authors:** Yasmine Adimi, Clement Levin, Emilia Puig Lombardi, Benjamin Morin, Claire Gervais, Sandrine Tolou, Anne-Claude Noirot-Jagu, Julie Hauret, Corinne Thomas, Patricia Tchepikoff, Fedor Grigorev, Anya Djafri, Sandrine Longuemaux, Anne Caron, Souad Naimi, Matteo Cesaroni, Eric Tartour, Marion Classe, Cecile Badoual

**Affiliations:** Paris Cité University, INSERM U970, PARCC, Paris, France; Sanofi PMCB, Research and Development department, Vitry, France; Pathology department, Hôpital Européen Georges Pompidou, APHP, Paris, France; Oncology department, Hôpital Européen Georges Pompidou, APHP, Paris, France

## Abstract

In the ever-changing world of digital pathology, being able to extract a maximum amount of information from a patient tissue sample is of paramount importance for better diagnosis, disease characterization, and therapeutic strategies. Recent technologies such as multiplex immunofluorescence imaging and spatial transcriptomic now enable a deep analysis of protein and gene expression while retaining the spatial context of the tissue. Here, we describe an innovative approach combining a 34-protein Phenocycler panel and transcriptome analysis using Visium on a single head and neck squamous cell carcinoma section. While protein analysis reveals the complexity of the immune phenotypes involved in the disease, transcriptome analysis reveals the intricate cellular states of cancer cells that coexist within the patient’s tumor. Finally, integrating both omics modalities, we uncover unique comparison of gene and protein expression of spatially resolved cellular subspaces.

## Introduction

Head and neck squamous cell carcinomas (HNSCC), affecting areas such as the oral cavity, pharynx, and larynx, are significant health concerns globally due to their high rates of illness and death^1,2^. These cancers arise from various causes, with the primary risks being tobacco use, alcohol consumption, and infection with high-risk human papillomaviruses (HPV), particularly HPV16^3–5^.Treatment options for HNSCC are diverse and can include surgery, radiation, chemotherapy, and increasingly, targeted therapies and immunotherapy, tailored to the tumor’s location and stage^2,6–9^. HNSCC ranks as the sixth most common type of cancer worldwide^2^. Survival rates vary significantly, mostly depending on the cancer’s stage and HPV status at diagnosis. Patients with HPV-positive HNSCC tend to have better outcomes, with a five-year survival rate of 85-90% for localized disease, substantially higher than the 55-60% survival rate for those with HPV-negative cancers^10–14^. This improved prognosis is partly due to HPV-positive tumors responding better to treatments such as radiation and chemotherapy^15^. Additionally, recent studies showed that these patients also benefit from immune checkpoint inhibitors, likely related to the unique immune characteristics of HPV-related tumors^16^. This difference highlights the importance of HPV status as a critical prognostic factor and underscores the necessity for precise diagnostic tools and personalized therapeutic approaches to optimize patient outcomes.

Indeed, Recent advances in omics technologies have profoundly transformed cancer research, enabling unprecedented molecular characterizations and the identification of biomarkers that are crucial for predicting therapeutic responses and facilitating early detection^17^. Spatial omics technologies, which integrate high-throughput molecular data within the precise spatial context of tissue architecture, provide comprehensive insights into the molecular heterogeneity present within tumors. This integration is essential for developing personalized treatment strategies tailored to the unique molecular profiles of individual tumors.

Among the standout spatial omics platforms, 10X Genomics Visium and Akoya PhenoCycler-Fusion® (ex-CO-Detection by indEXing - Codex) are particularly noteworthy for their innovative approaches to capturing molecular data^18^. Visium, developed for spatially resolved transcriptomics, captures the entire transcriptome within its histological context, allowing researchers to observe gene activity across the entire landscape of a tumor^17,19^. This broad view is invaluable for understanding how different regions within a tumor may respond to various treatments or how they might contribute to disease progression. Conversely, Phenocycler offers a complementary approach through high-resolution proteomic imaging. It can map multiple proteins simultaneously at the cellular level, providing detailed insights into active biological processes^17,20,21^. This capability is crucial for a more nuanced understanding of cellular functions and the interactions within the tumor microenvironment that are not visible through transcriptomic data alone.

Phenocycler’s technology is particularly revolutionary due to its ability to analyze more than 50 markers on a single tissue slide using a cyclic multiplexing method^17,21,22^. This process involves applying antibodies and sets of fluorescently tagged oligonucleotide probes to a tissue section, imaging them, and then chemically removing the tags to prepare the section for the next set of probes^20^. This cycle can be repeated multiple times, accumulating a vast dataset from the same tissue section without damaging it^20^. This extensive profiling allows researchers to dissect complex cellular phenotypes and elucidate intricate signaling pathways, offering significant insights into potential therapeutic targets.

Visium and Phenocycler together provide a powerful and complementary combination for cancer research. While Visium gives a comprehensive snapshot of gene expression patterns, Phenocycler adds a layer of proteomic detail that is vital for understanding the functional state of proteins within the cells. This combination allows for a multi-dimensional view of both RNA and protein expressions, significantly enhancing the ability to discern subtle yet critical variations within the tumor microenvironment that may influence disease progression and response to treatment. Thus, the integration of these technologies marks a significant advancement in the field of precision oncology.

Current approaches for analyzing spatial omics often face significant challenges due to the integration of data derived from separate tissue slides for proteomic and transcriptomic assays, such as those generated with Phenocycler and Visium, respectively^23,24^. This separation between modalities can lead to the loss of crucial information, as the cellular structures and spatial organization may not be consistent between different sections. Consequently, correlating protein and RNA data from these disparate sources can result in discrepancies, limiting our ability to draw precise conclusions about the cellular mechanisms at play within the tumor microenvironment.

Despite their profound impact, each technology has limitations when used independently. Phenocycler, although precise in protein localization, is limited by the number of targets that can be analyzed, potentially overlooking significant pathways active within the tumor microenvironment.

A fundamental limitation of Visium lies in its spot-based resolution, which, while providing a broad overview of gene expression across a tissue section, may miss finer details at the cellular level. This resolution gap means that Visium can occasionally overlook critical spatial relationships and cellular interactions that are vital for understanding complex biological processes.

In an attempt to address these challenges, our study focuses on integrating Phenocycler directly on the same tissue slide used for Visium, providing a promising approach to overcoming these obstacles.

In this study, we applied this innovative spatial omics approach on an initial diagnostic biopsy from a 61-year-old patient diagnosed with HPV-positive left tonsillar cancer, staged as TNM T4N2M0. Initially treated with radiotherapy, the patient experienced a recurrence at the primary site and regional lymph nodes within eight months. However, the disease subsequently metastasized to the lungs, pleura, liver, and hilar regions. Treatment with nivolumab did not elicit a response, and the cancer progressed. By employing both proteomic and transcriptomic profiling on a single slide of the biopsy of this patient, we can achieve a more comprehensive view, capturing both RNA and protein expressions within the exact same histological context. To ensure reproducibility, we conducted both proteomic and transcriptomic profiling on two different sections of the same biopsy using Phenocycler and Visium platforms. This co-localization not only ensures that the structural integrity and cellular context are preserved but also enhances data richness, allowing for a more accurate comparison and correlation between protein and RNA data. Additionally, we applied the same approach to a separate sample to further validate our findings.

### Results

### 1. Validation of the Integrated Phenocycler and Visium Assay on a Single Slide

To comprehensively assess the spatial distribution of immune cells and their transcriptomic profiles within the same sample, we employed a dual approach of spatial proteomics (Phenocycler) and spatial transcriptomics (10x Visium) on the same HNSCC slide and tried to investigate their collective impact on RNA integrity and gene expression in head and neck squamous cell carcinoma (HNSCC) samples. Figure 1a. illustrates our experimental design, which compares a sample processed by both proteomics and transcriptomics (PP+ST) with a consecutive section analyzed solely by transcriptomics (ST only).

**Fig. 1.**
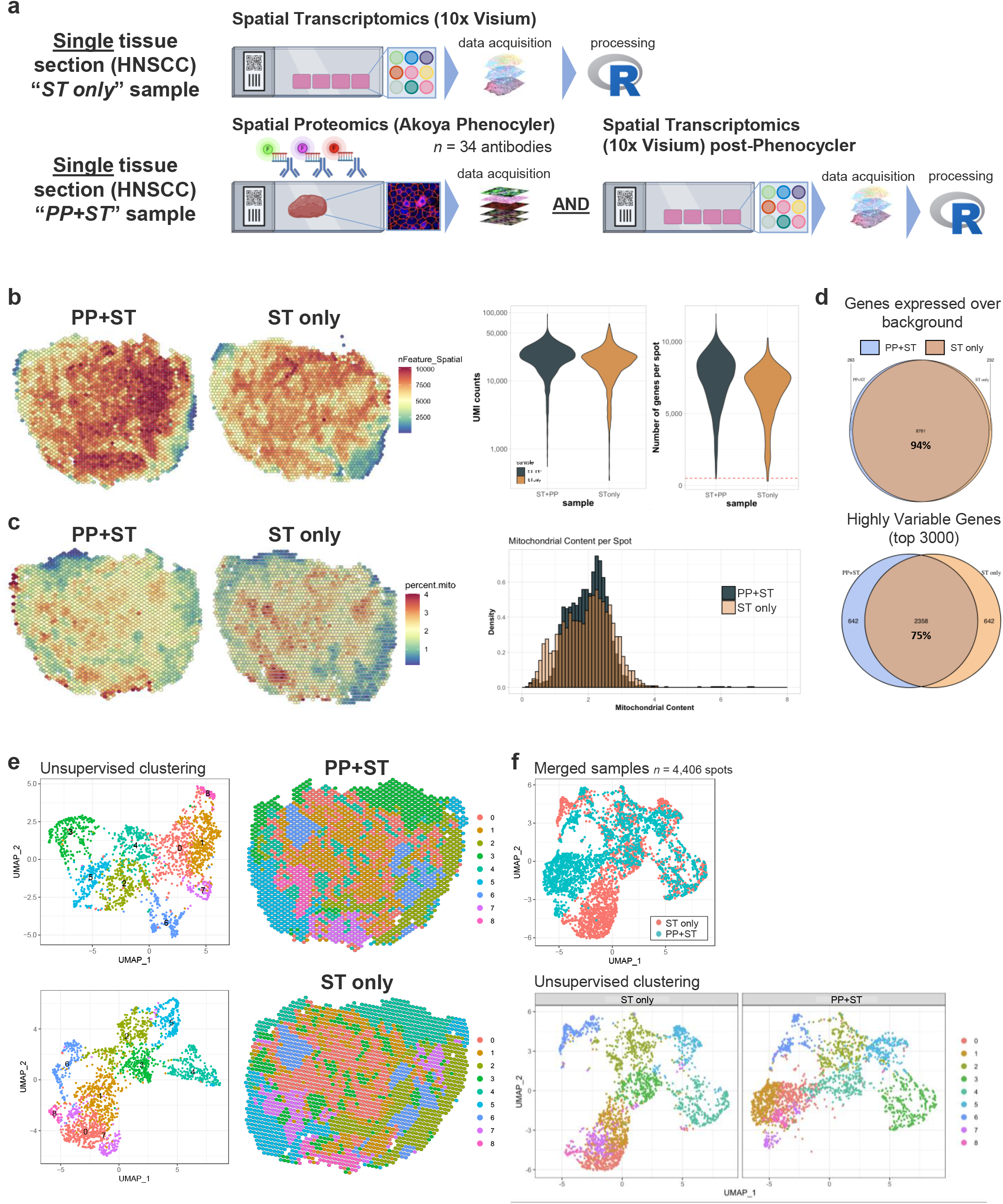
Integrated spatial proteomics and transcriptomics in head and neck squamous cell carcinoma (HNSCC). (a) Workflow for “ST only” and “PP+ST” samples using spatial transcriptomics and spatial proteomics. (b) Spatial distribution of gene (feature) and UMI detection per spot in *PP+ST* and *ST only* samples. For violin plots, the y-axis is set on a log_10_ scale for clarity. (c) Mitochondrial content comparison between *PP+ST* and *ST only* samples. (d) Venn diagrams showing the overlap of genes expressed above background (top) and of the top 3000 highly variable genes (bottom) detected in *PP+ST* and *ST only* samples. (e) Transcriptomic landscape patterns derived from unsupervised clustering in *PP+ST* and *ST only* samples. (f) UMAP projection of 4,406 spots after merging of the *PP+ST* and *ST only* samples.

In our methodology for the PP+ST sample, we first conducted Phenocycler analysis followed by Visium. This sequence was crucial because the Visium protocol involves tissue permeabilization that could potentially alter cellular structures, thereby affecting the resolution necessary for effective Phenocycler analysis. Therefore, we did not start with Visium to avoid compromising the structural integrity required for subsequent Phenocycler imaging. Overall, after filtering low-quality spots and cells (see *Methods*), we captured similar numbers of spots for analysis in both conditions, 2,202 and 2,177 spots in the PP+ST and ST only samples, respectively.

Contrary to initial concerns about RNA degradation, our findings indicate that RNA integrity in the PP+ST sample was maintained comparably to the ST only samples, as demonstrated by the similar distributions of total gene expression (Fig. 1b). Remarkably, the PP+ST sample exhibits a dense concentration of high-feature spots throughout the tissue section, suggesting that the proteomic pre-treatment does not compromise the detection capacity of the subsequent transcriptomic analysis.

Another relevant quality control metric is the quantification of unique molecular identifiers (UMIs) paired with the evaluation of the number of detected genes per spot. While the UMIs show a comparable distribution of counts between PP+ST and ST only, maintaining high levels of transcript detection, the plots of the number of genes per spot reveal a slight shift toward higher gene counts in the PP+ST sample (Fig. 1b), aligning with the noticeable increase in mitochondrial transcript density observed in the same sample (Fig. 1c), which suggests a potential higher presence of damaged cells in this sample. This increase in mitochondrial transcripts contributes to the overall higher gene counts compared to the ST only samples.

Furthermore, analysis of gene expression levels shows that 94% of genes expressed above background signal are common to both the PP+ST and ST only samples, indicating robust preservation of gene expression across both experimental setups. Additionally, the Venn diagram depicting the top 3000 highly variable genes (HVGs) for each sample shows that 75% of these genes are shared between the two methods, with each condition contributing unique genes to the remainder (Fig. 1d). The differences observed for HVGs between the PP+ST and ST only samples may be explained by the use of consecutive tissue sections for each condition. Despite the differences in HVGs detected in the PP+ST and ST only samples, the results of unsupervised clustering represented through UMAP visualizations highlight similar clustering patterns between PP+ST and ST only samples (Fig. 1e). This similarity underscores that, at a transcriptomic level, the overall cellular heterogeneity and molecular signatures are well-maintained despite the additional proteomic preprocessing.

Expanding on these findings, the merged UMAP plots allow for a direct comparison of spatial transcriptomic profiles, demonstrating a significant overlap in the data points from both PP+ST and ST only samples (Fig. 1f). with the exception of some spots that we later annotated as malignant cells. Given that the sampling is not at a precise cellular scale, slight variations in cellular composition between sections can lead to differences in gene expression profiles, particularly in spots covering tumor cells (see Discussion).

To further ensure reproducibility and robustness of our findings, we performed similar dual analyses (PP+ST and ST only) on two separate samples from the same patient, as well as on an additional sample from a different patient. These additional experiments (Extended Data Figs. Supp.1 and. Supp.2), confirm the consistency of our observations across different samples, thereby reinforcing the validity of our integrated approach.

In all, this analysis confirms the feasibility and efficacy of integrating spatial proteomics with transcriptomics on the same tissue section, ensuring high fidelity in molecular profiling, which is crucial for advancing research.

### 2. Intratumoral Diversity and Cellular Dynamics in HNSCC Revealed Through Multiplex Immunofluorescence

We developed a 34-antibody multiplex immunofluorescence (mIF) Phenocycler panel to characterize the tumor microenvironment of HNSCC. Using this combination of markers, we are able to identify various cell types and understand their functions. The panel includes antibodies for immune cell phenotypes such as CD38, CD3e, CD4; structural proteins like Vimentin and aSMA; checkpoint molecules such as PD1 and LAG3; and other cell status markers such as Ki67 for proliferation (Fig. 2a).

**Fig. 2.**
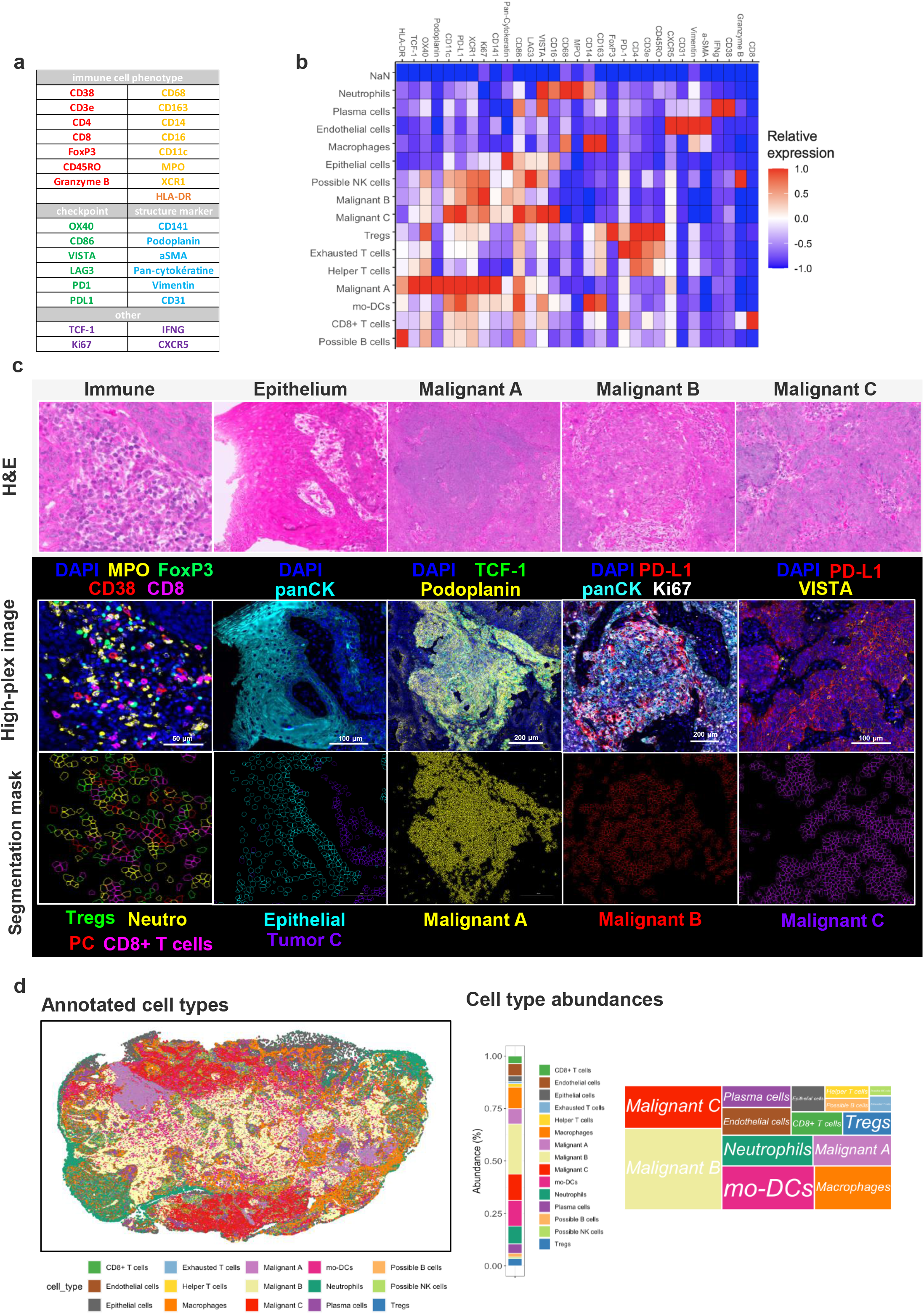
Intratumoral diversity in HNSCC revealed through multi-immunostaining. (a) Antibody panel for Phenocycler analysis categorized by their target groups: immune cell phenotypes, structural markers, checkpoint inhibitors, and other relevant markers. (b) Heatmap of relative marker expression in the different annotated cell types (c) Top: Selected regions of interest from H&E staining depicting immune cells, epithelial cells, and the three malignant clusters for histological context. Middle: multiplex immunofluorescence images from same regions of interest, revealing heterogeneity of protein marker expression. Bottom: Segmentation masks of clusters. (d) Left: Overlay of annotated cell types on the HNSCC tissue image. Right: Bar chart depicting the abundance of each cell type.

After image QC and cell segmentation using the Deepcell algorithm, successfully distinguishing and capturing a diverse array of cellular morphologies without the need for manual corrections or annotations, unsupervised clustering was performed and sixteen distinct cell clusters within the tumor microenvironment were found, including three malignant clusters—Malignant A, Malignant B, and Malignant C—each marked by unique proteomic signatures (Fig. 2b). Malignant A is characterized by densely packed cells indicative of aggressive growth, while Malignant B and C feature cells that are more sparsely distributed, indicating a less compact cell structure. Fluorescence imaging distinguishes these clusters further: Malignant A shows high expression of pan-cytokeratin and TCF-1, indicating robust epithelial origin and proliferative activity; Malignant B is marked by significant PD-L1 expression, suggesting adaptive immune resistance; and Malignant C exhibits prominent VISTA expression, associated with immune suppression. Segmentation maps detail the spatial context, highlighting interactions with surrounding immune cells such as Tregs and CD8+ T cells, which might influence the immune environment of each cluster. This detailed visualization provides crucial insights into the complex tumor heterogeneity within HNSCC (Figs. 2b, c). Unsupervised clustering also reveals a rich and diverse immune cell environment, populated by plasma cells (PC), neutrophils, T cells, macrophages, among others. The segmentation mask highlights the physical proximity and potential interactions among these different immune cell types within the tumor stroma, indicating a complex immune landscape that could influence tumor behavior and response to treatments (Figure 2c).

Mapping of phenotypic clusters on the sample image reveals a structured spatial organization of malignant and immune cell types, arranged in well-defined areas of the tissue (Fig. 2d). The spatial arrangement of the malignant clusters might indicate areas of varying tumor aggressiveness and microenvironmental influence, with implications for tumor growth, metastatic potential, and response to therapy.

A quantitative bar chart details the proportion of each cell type, with significant presence of immune cells such as Tregs and neutrophils. This visualization highlights the heterogeneity and the intricate interactions between tumor and immune cells in HNSCC (Fig. 2d).

### 3. Integrated Multi-Omic Profiling Reveals the Cellular and Molecular Dynamics in Head and Neck Squamous Cell Carcinoma (HNSCC)

Having comprehensively annotated the slide on a true single-cell protein level and given that our transcritomics assay suffers from low spatial resolution, we sought to transfer the information assayed by Phenocycler onto the Visium modality. The integration of the spatial transcriptomics and proteomics modalities on the same tissue slide involved the meticulous alignment of the two datasets, where the Visium data provided a spatial grid of gene expression that was overlaid with the high-resolution Phenocycler image and segmentation masks to precisely delineate cellular boundaries (see *Methods*). During the image registration step, we scaled and shifted the Phenocycler image to perfectly match the Visium coordinates, ensuring accurate association of proteomic measurements with corresponding transcriptomic data points. This alignment facilitated the aggregation of cell-level measurements from Phenocycler into the pseudo-spot level of the Visium grid, thereby creating a unified dataset that captures both proteomic and transcriptomic profiles of distinct microenvironments within the HNSCC sample (Fig. 3a). After conversion of single-cell protein expression into pseudospot resolution, consistent patterns and levels of expression are observed for selected markers (Fig. 3b).

**Fig. 3.**
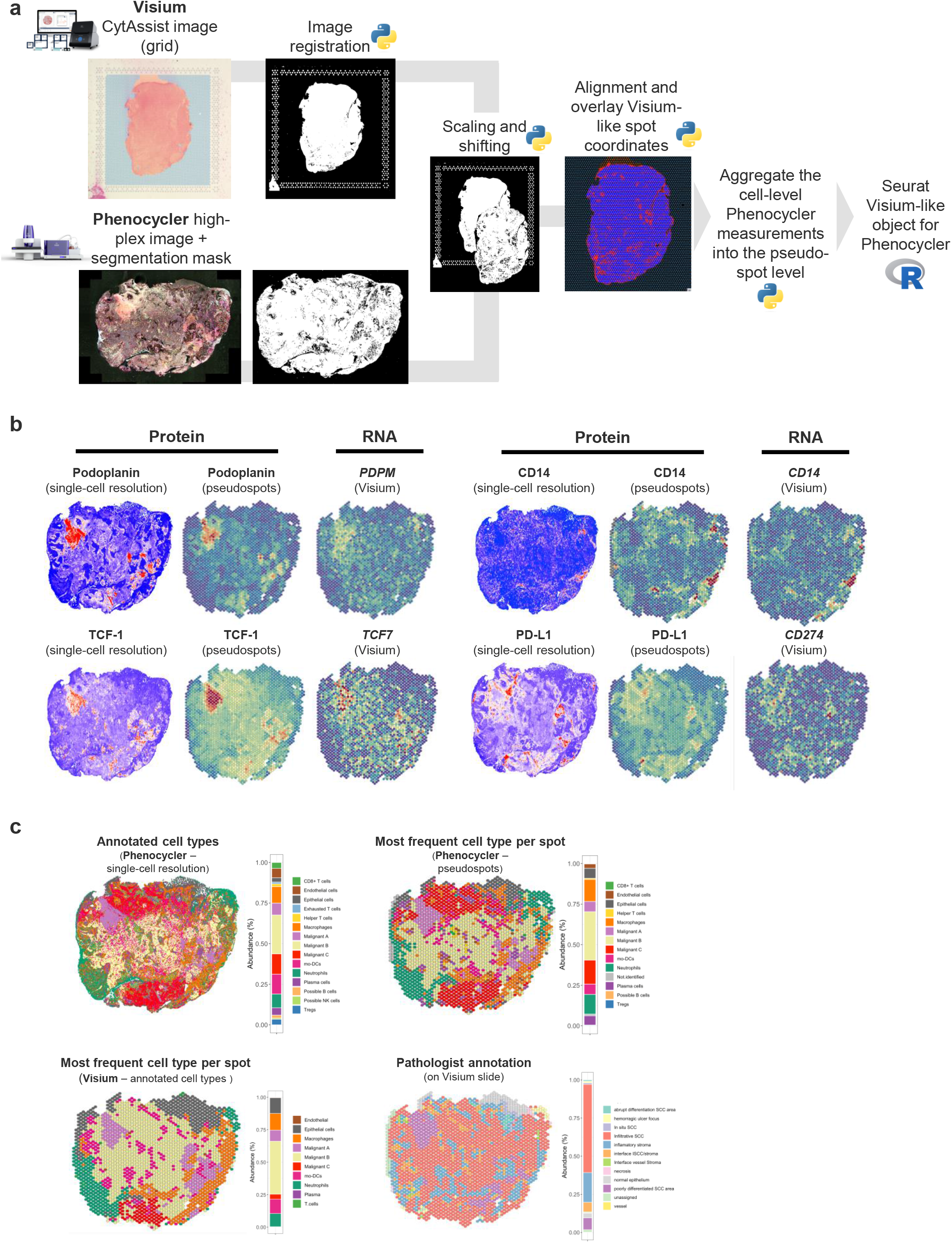
Integrated multi-omics profiling reveals the cellular and molecular dynamics in HNSCC. (a) Workflow for integrating spatial transcriptomics (Visium) and spatial proteomics (PhenoCycler) on a single tissue section. (b) Side-by-side comparison of single-cell and pseudospot resolutions for protein markers Podoplanin, PD-L1, CD14, and TCF-1 from PhenoCycler, alongside Visium gene expression data. (c) Cell type annotations from PhenoCycler at single-cell resolution; most frequent cell types per pseudospot from PhenoCycler; annotated cell types from Visium transcriptomic data, indicating cellular composition and spatial distribution.

Furthermore, both Phenocycler pseudospots and Visium data showed similar spatial distributions for Podoplanin, TCF1 (encoded by *TCF7* gene) and CD14, indicating a strong concordance between protein and gene expression within these regions. However, the expression patterns for PDL1 (encoded by *CD274* gene) demonstrated notable discrepancies; Phenocycler data revealed more localized and intense expression which was not as prominently mirrored in the Visium data, suggesting potential differences in protein activity versus mRNA presence (Fig. 3b). This difference underscores a critical limitation of the Visium technology, particularly in capturing rare or spatially scattered cell types. The Visium technology, primarily designed for broader transcriptomic surveying, may not adequately reflect the nuanced, localized expression patterns that are crucial for understanding cellular function in a heterogeneous tissue landscape. Consequently, the integration of Phenocycler data provides a more detailed and localized view of protein activity that complements and enhances the broader gene expression profiles captured by Visium.

Consistent clustering results were also observed between single-cell and pseudospot resolutions using Phenocycler data, confirming similar cellular distributions and marker expressions across both scales. The annotations derived from the Phenocycler data were then applied to annotate clusters identified in the Visium transcriptomic data. Finally, we compared the annotated cell types between Phenocycler pseudospots and Visium, finding significant correlations in the spatial distribution of key cell types, such as the malignant clusters and immune cells, across both datasets. This cross-validation not only underscored the robustness of our clustering approaches but also confirmed the presence and the distribution patterns of critical cell populations within HNSCC, as identified independently by proteomic and transcriptomic profiling (right panel, Fig. 3c). However, populations such as Tregs and B cells, which are challenging to distinguish in Visium due to their typically sparse distribution, were rarely dominating in any single Visium spot. This is a limitation in Visium’s resolution, as it often averages out the cell types within a larger, predefined spot, potentially obscuring less abundant but biologically significant cells. In contrast, Phenocycler offered a more granular view with enhanced precision, allowing for detailed identification and quantification of these rarer immune cells at specific locations. This increased resolution provided by Phenocycler was crucial, as it brought much-needed clarity to the cellular composition within the tumor microenvironment, highlighting the presence of specific immune cells that Visium could not distinctly resolve. Thus, while both technologies showed overall correlations in broader cell type distributions, Phenocycler’s precision significantly enriched our understanding of the immunological aspects within the tumor landscape. This comparative analysis not only underscores the complementary nature of the two platforms but also emphasizes the need for integrating such detailed proteomic data to gain a complete picture of the cellular dynamics in cancer especially since, the pathologist’s annotations, which emphasize visible and dominant morphological features such as tumor morphology and primary cell type distributions, showed a general alignment with the broader, spot-based patterns depicted in the Visium data (Fig. 3c).

In conclusion, we successfully integrated spatial transcriptomics (Visium) with spatial proteomics (Phenocycler) on the same tissue slide, significantly enhancing our understanding of the tumor microenvironment with detailed resolution.

### 4. Differential Gene Expression and Pathway Analysis accros the different malignant states

The identification of three malignant cell clusters in both Phenocycler and Visium datasets directed our focus towards these groups, especially since Malignant state A shows notable similarity across both platforms. This prompted a detailed examination of these clusters to explore their characteristics (Fig.4a). Malignant A is marked by high levels of *SOX9* (Fig. 4c), *TCF7, Cd70* and *PDPN* (Figs. 4b, c). Malignant B features *FOXQ1, KREMEN1, MUC16* and *CLDN4* expression and Malignant C expresses elevated *ATF5, NURP1, VEGFA* and *EGLN3* (Figure 4b). These expression differences are likely linked to distinct levels of tumor cell differentiation, reflecting the complex heterogeneity and progression within the tumor.

**Fig. 4.**
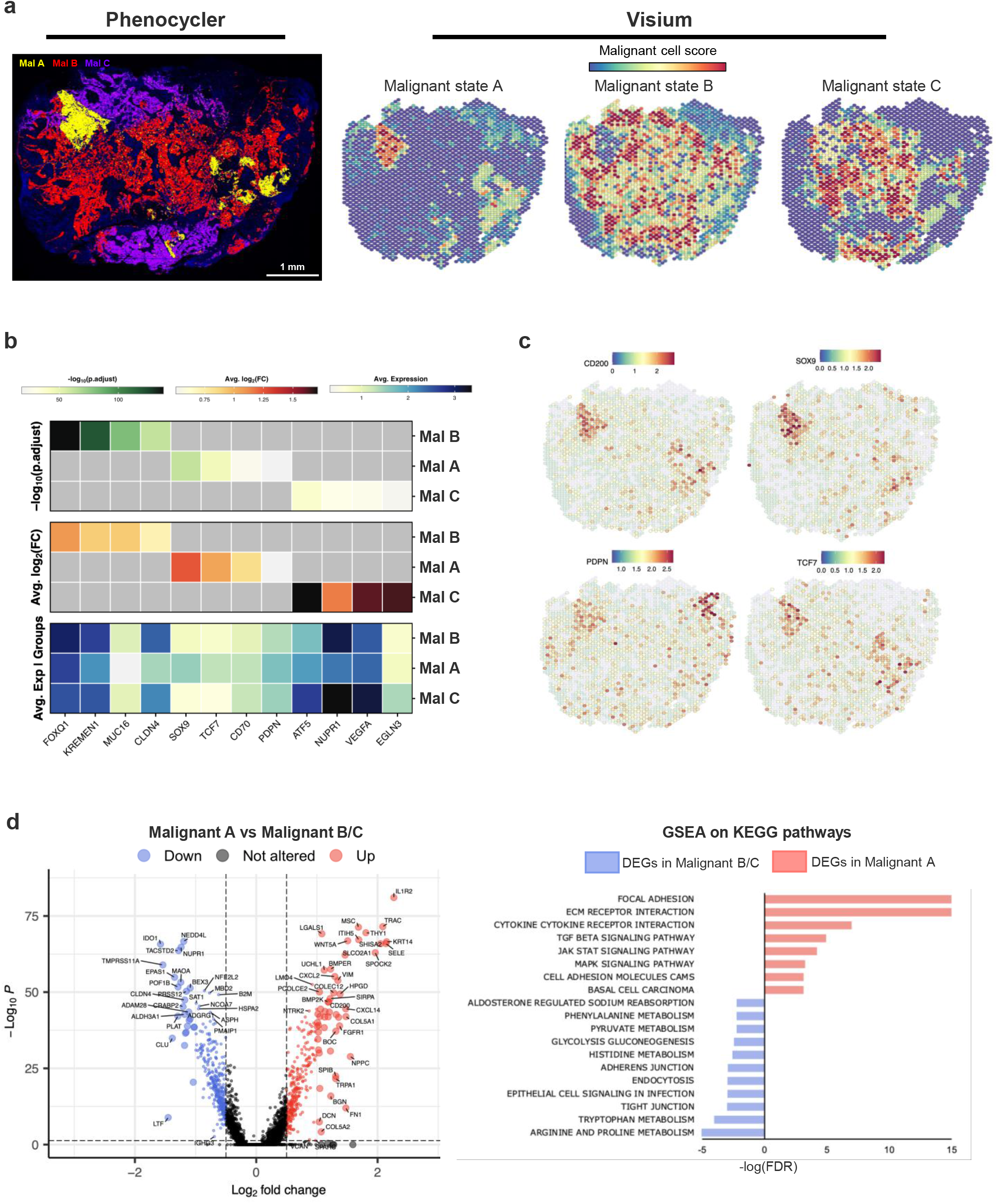
Molecular profiles of the malignant states. (a) Comparative imaging from Phenocycler and Visium platforms depicting three distinct malignant states (A, B, C) (b) Heatmap illustrating differential expression levels of key biomarkers across Malignant states A, B, and C.(c) Spatial distribution maps of selected biomarkers (*CD200, SOX9, PDPN, TCF7*) in tissue sections, showing the expression levels and localization of each marker in the tumor microenvironment. (d) Left: Volcano plot contrasting gene expression in Malignant A versus Malignant B/C, with upregulated genes in red, downregulated in blue, and unchanged in black. Right: Gene Set Enrichment Analysis (GSEA) highlighting key enriched pathways in Malignant A (red) and Malignant B/C (blue) based on differential gene expression.

Our analysis specifically focused on Malignant A due to its notable colocalization with Phenocycler results, indicating a robust alignment of transcriptional and proteomic data. To understand the unique biological features of Malignant A, we conducted a comparative analysis of expression patterns within these spots against the combined characteristics of Malignant B and C. In this cluster, genes such as *IDO1, ALDHLA3*, and *CLDN4* are significantly downregulated, indicating potential reductions in immune interaction and cellular adhesion. On the other hand, genes like *BMP2* and *CSCL12* are markedly upregulated. *BMP2*, pivotal in the TGF-beta signaling pathway. *CXCL12*, enhancing cytokine-cytokine receptor interactions.

Gene Set Enrichment Analysis (GSEA) further underscores these findings, with Malignant A showing substantial activation of pathways such as TGF-beta and JAK-STAT signaling, fundamental for cell signaling and growth regulation (Fig. 4d). Conversely, Malignant B and C display enrichment in pathways critical for structural integrity and tumor spread, such as focal adhesion and extracellular matrix (ECM) receptor interactions. These pathways are vital for maintaining cellular cohesion and facilitating migration (Fig. 4d).

To validate the elevated *SOX9* levels observed in Malignant A at the RNA level, we performed immunohistochemical staining to confirm SOX9 protein expression (Figs 5a). Malignant state A showed significantly higher levels of SOX9 expression compared to Malignant states B and C, validating transcriptomic data (Figs 5b and c). We analyzed the impact of *Sox9* expression on overall survival (OS) and progression-free survival (PFS) in patients with head and neck squamous cell carcinoma (HNSCC) from the TCGA database. Kaplan-Meier survival curves show that high *Sox9* expression is significantly associated with reduced overall survival (p = 0.037; Fig. 5d). Similarly, high *Sox9* expression correlates with significantly decreased progression-free survival (p = 0.033; Fig. 5e).

**Fig. 5.**
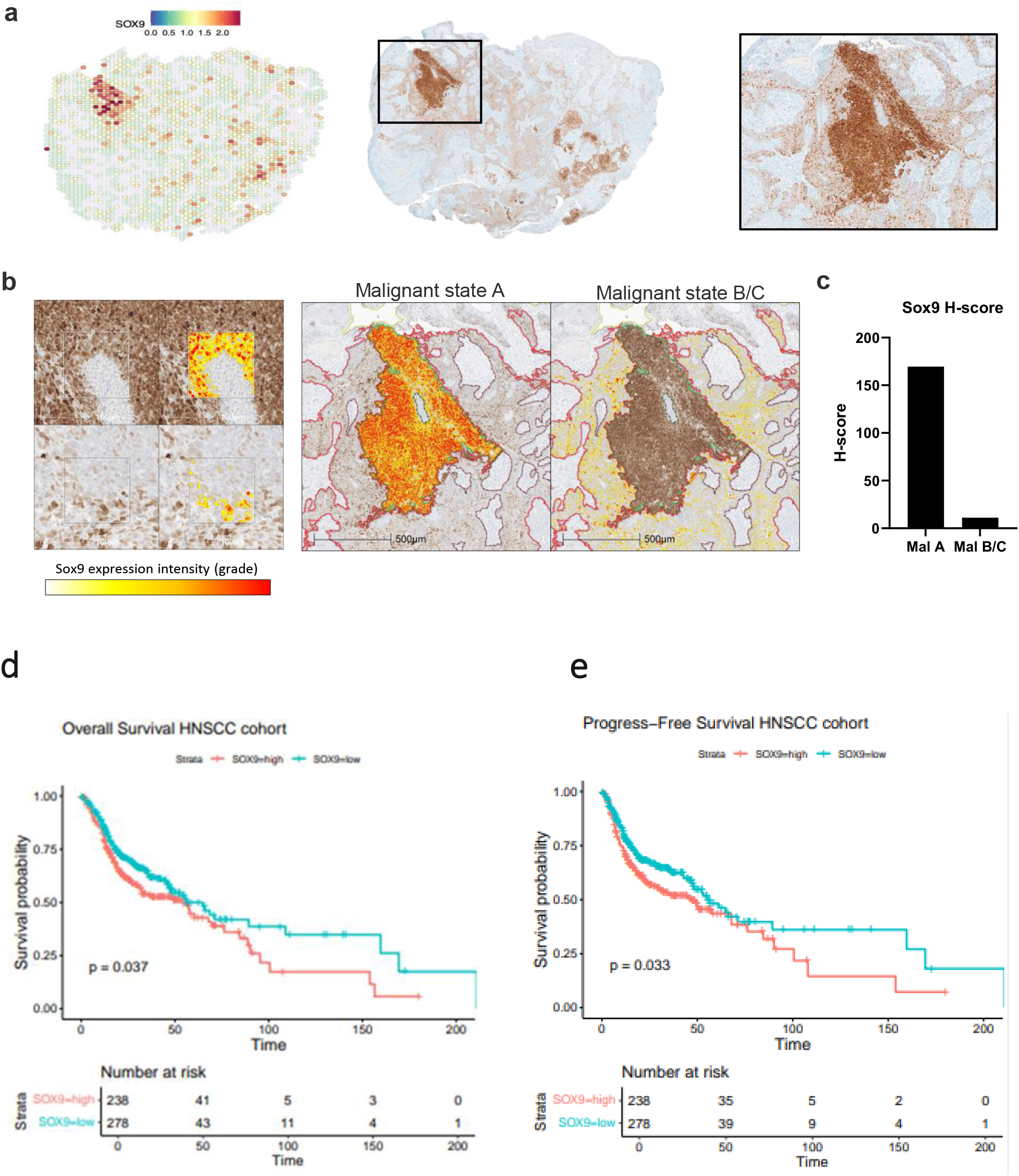
Validation of SOX9 expression by IHC and its survival impact in TCGA HNSCC cohort. (a) Spatial distribution of *SOX9* expression from Visium data and corresponding immunohistochemical staining using an antibody targeting SOX9 protein. (b) Detailed comparison of SOX9 expression intensity in Malignant state A and Malignant state B/C. (c) H-score quantification of SOX9 expression between Malignant state A and Malignant state B/C. (d) Overall Survival of TCGA-HNSCC patients grouped by *SOX9* expression levels. (e) Progression free Survival of TCGA-HNSCC patients grouped by *SOX9* expression levels.

In addition to SOX9, we also investigated the expression of CD70 in Malignant state A. Immunohistochemical staining revealed visibly higher CD70 protein expression in Malignant state A compared to states B and C, consistent with the transcriptomic data (Extended Data Fig. S3).

## Discussion

In this study, we combined spatial transcriptomics and proteomics data from a single HNSCC histological slide, achieving an unprecedentedly detailed understanding of the tumor microenvironment. In general, understanding the tumor microenvironment (TME) is fundamental in cancer research due to its profound influence on tumor development, progression, and therapeutic response. The TME comprises a complex network of cancer cells, immune cells, stromal cells, blood vessels, and extracellular matrix components ^25^. This intricate ecosystem not only supports tumor growth and metastasis but also plays a critical role in immune evasion and resistance to treatment ^26^. Advanced techniques for studying the TME, including high-dimensional single-cell analysis with mass cytometry (CyTOF), single-cell RNA sequencing (scRNA-seq), bulk RNA sequencing, offers detailed phenotypic and transcriptomic data but lacks spatial context ^27^. Spatial transcriptomics, and multiplex imaging, provide diverse and complementary insights. These methods enable a detailed characterization of cellular phenotypes, interactions, and spatial organization within the tumor tissue ^27,28^. In contrast, spatial techniques retain spatial information and provide insights into the organization and interactions within TME ^27^. Spatial transcriptomics methods, such as MERFISH (Multiplexed Error-Robust Fluorescence In Situ Hybridization) and 10x Genomics Visium, enable the localization of gene expression within tissue sections, preserving spatial context ^28,29^. These techniques provide high-throughput spatial mapping of gene expression, crucial for understanding the spatial heterogeneity of tumors. Multiplex imaging methods, such as Imaging Mass Cytometry (IMC), Multiplexed Ion Beam Imaging (MIBI), and CODEX (CO-Detection by Indexing), allow for the visualization of multiple proteins simultaneously within tissue sections, retaining spatial information and revealing complex cellular interactions. In situ hybridization (ISH) techniques, like RNA-ISH and DNA-ISH, detect specific nucleic acid sequences within fixed tissues, preserving spatial context ^28,29^.

In recent years, several studies have aimed to connect RNA and protein expression while preserving spatial information ^23,24^. To address these questions, techniques like Phenocycler and Visium are usually applied to consecutive slides ^23,24^. However, this consecutive application can introduce variability due to differences in tissue integrity and cellular composition. Indeed, slight variations between sections can significantly impact gene expression profiles (Fig. 1d), especially in tumor areas, highlighting the challenges of analyzing adjacent but separate tissue sections. Additionally, aligning images from consecutive slides is technically challenging, whereas working on images from the same histologic section, as shown by our method, is straightforward.

We specifically chose to sequence the Phenocycler analysis first, followed by Visium, because Phenocycler has the distinct advantage of preserving tissue integrity after processing^20^. Unlike other methods that may denature the tissue and obscure certain details, Phenocycler maintains the structural integrity of the sample ^20^. Additionally, Phenocycler utilizes a flow cell that enables comprehensive imaging of the entire sample, ensuring that no regions are missed during analysis. This approach contrasts with other histological methods that can degrade tissue quality and add troublesome complexity to the workflow.

Therefore, our integrated approach preserves the spatial context and continuity of the tissue architecture, ensuring more accurate and reliable data correlation and interpretation. The integration of these multiomics data allowed us to identify three distinct clusters of malignant cells with unique gene and protein expression profiles (Fig. 3c). Notably, Cluster A showed a strong correlation between molecular data, proteomics data, and the pathologist’s annotations, providing deeper insights into its specific tumor biology. Cluster A exhibited significant upregulation of the cell cycle control and cellular differentiation genes *SOX9* and *TCF7*, confirmed at the protein level.

SOX9 plays a crucial role in maintaining stem cell properties and is essential for the differentiation of various cell types ^30^. It has also been linked with cancer progression and metastasis, influencing genes involved in cell proliferation, survival, and invasion ^31,32^. Overexpressing SOX9 is linked to increased tumor-initiating capabilities and resistance to conventional therapies, making it a marker of cancer stem cells and a potential therapeutic target ^31,32^. Significantly, high SOX9 levels in poorly differentiated malignant cluster suggest these cancer cells maintain a stem-like, undifferentiated state, which is associated with higher tumorigenic potential, resistance to conventional treatments.

On the other hand, *TCF7*, which encodes TCF1, is crucial for T-cell lineage commitment and maintaining a balance between proliferation and differentiation in cancer ^33,34^. It is part of the Wnt signaling pathway, regulating cell fate, proliferation, and migration. In cancer, TCF1 helps maintain the undifferentiated state of cancer cells, contributing to tumor growth and playing a role in the tumor immune environment, potentially aiding in immune evasion ^35^.

Interestingly, our analysis also identified elevated levels of CD70 in Malignant state A. CD70, a molecule induced by TGF-β, is often associated with epithelial-mesenchymal transition (EMT), a key process in cancer progression and metastasis^36^. The presence of CD70 in Malignant state A suggests an active EMT process, which could contribute to the aggressive nature of these cells. EMT is known to promote cellular plasticity, enabling cancer cells to acquire invasive and metastatic capabilities. This association with EMT further supports the identification of Malignant state A as a poorly differentiated and highly aggressive tumor cell population.

The pathologist’s classification of Cluster A as “poorly differentiated” aligns with our multi-omics findings, highlighting these cells’ aggressive nature. Poorly differentiated tumors typically grow rapidly and spread quickly. The molecular profile of Cluster A, with high levels of SOX9, TCF1, and CD70, supports this, offering insights into what drives its aggressive behavior. The proteomic data also highlighted the functional implications of these gene expression patterns. High SOX9 levels suggest active signaling pathways that promote stemness and metastasis, while increased TCF1 levels indicate enhanced proliferation and immune modulation. Additionally, the upregulation of CD70, linked to EMT, underscores the plasticity and invasive potential of these cancer cells. This combination of traits makes Cluster A a highly adaptable and resilient cancer cell population, capable of sustaining tumor growth and evading the immune system, which can explain the patient’s non-responsiveness to immunotherapy.

Furthermore, each spatial omics modality, Phenocycler and Visium, provided unique insights into the tumor microenvironment, showcasing the strengths of each approach and the enhanced value of their combination. Phenocycler delivered high-resolution proteomic data, allowing detailed visualization of protein distributions within cells. Unlike transcriptomic data, proteomic data directly reflects cellular function, offering an immediate and functional view of cellular processes. Additionally, our use of a tailored panel of 34 markers enabled precise identification and characterization of cellular heterogeneity within the tumor, identifying three distinct malignant clusters and allowing annotation of 15 different cell types, each defined by unique protein expression patterns.

Visium complemented Phenocycler by providing a broad transcriptomic overview of the entire tumor section. This spatial resolution is essential for correlating localized protein activities with corresponding transcriptomic data, offering a more complete picture of tumor biology and its microenvironmental interactions. The integration of Phenocycler and Visium data provided a synergistic approach that significantly enhanced our understanding of the tumor microenvironment. A key aspect of this integration was the creation of pseudospots from Phenocycler data, which were instrumental in achieving a more precise tissue annotation, aggregating high-resolution protein data into spatially coherent units that approximate the resolution of Visium spots. This method retains detailed protein expression information while enabling direct comparison with Visium’s transcriptomic data. Using pseudospots, we refined our cellular annotations beyond the broader categories typically identified through Visium’s transcriptomic deconvolution.

However, the current methodology of transitioning from the cellular scale in Phenocycler data to the spot scale of Visium to achieve concordance between the datasets introduces a significant bias in our analysis. This approach inherently dilutes the high-resolution data provided by Phenocycler, as aggregating individual cell data into larger spot-based clusters can obscure specific cellular details and interactions that are critical for understanding microenvironmental dynamics. Ideally, the reverse process would be more beneficial, expanding the lower resolution Visium data to match the detailed cellular resolution of Phenocycler. Currently, Visium technology maps transcriptomic data to predefined spots rather than to individual cells, which can obscure finer details of cellular heterogeneity and microenvironmental interactions, as each spot may contain multiple cell types, making it challenging to attribute specific gene expressions to distinct cellular behaviors.

Looking forward, advancements such as Visium HD or the forthcoming Xenium® platform promise to address these limitations, offering higher resolution at or near the single-cell level, significantly improving our ability to dissect complex biological processes within tumors.

## Material and methods

### Tissue material

Sample was obtained from a diagnosis biopsy that had been embedded in an FFPE block at the European hospital Georges Pompidou, Paris, France. Patient provided written informed consent (CPP 2022-10-13).

### Phenocycler

A 5 *µ*m-thick section was produced from the FFPE block. The section was allowed to air-dry for 5 days. Subsequently, it was deparaffinized by placing it in an oven at 60 degrees Celsius overnight.

We conducted pretreatment using xylene and ethanol. Antigen retrieval was performed using AR9 buffer in a pressure cooker for 20 minutes. Subsequently, we incubated the tissue with antibodies in two steps, each lasting 3 hours at room temperature.

The specific antibodies used for tissue staining were anti-CD20-BX007, anti-CD21-BX032, anti-CD86-BX021, anti-CD11c-BX024, anti-CD38-BX089, anti-CD68-BX015, anti-CD163-BX069, anti-CD16-BX030, anti-VISTA-BX040, anti-CD14-BX037, anti-CD141-BX087,anti-CD3e-BX045, anti-CD4-BX003, anti-CD8-BX026, anti-FoxP3-BX027, anti-TCF-1-BX061, anti-Lag3-BX055, anti-CD45RO-BX017, anti-GranzymeB-BX041, anti-Ki67-BX047, anti-PD1-BX046, anti-PDL1-BX043, anti-OX40-BX029, anti-HLA-DR-BX033, anti-CXCR5-BX050, anti-IFNG-BX020, anti-Pan-cytokeratine-BX019, anti-Vimentin-BX022, anti-CD31-BX001, anti-Podoplanin-BX121, anti-XCR1-BX023, anti-aSMA-BX004, anti-MPO-BX098.

Probe addition, washing, and denaturing steps were executed using the PhenoCycler-Fusion instrument from Akoya Biosciences, version 2.1.0. The slide was then stored in a storage buffer for an additional 3 days before Visium processing.

### Pre-processing of Phenocycler data

In this study, the processing of QPTIFF images generated by the Phenocycler was conducted entirely on the Enable Medicine platform (https://www.enablemedicine.com/). Cell segmentation was performed using the DeepCell algorithm, a deep learning-based feature integrated within Enable Medicine, ensuring precise high-content image analysis. Following segmentation, rigorous quality control was implemented, with minor adjustments made to refine the acquired cellular outlines. Subsequently, unsupervised clustering was applied to group cells into phenotypically distinct clusters. These initial clustering results were carefully adjusted to better reflect the observed phenotypes. Preliminary annotations of these clusters were then verified by overlaying them onto the original images using Enable Medicine’s advanced visualization features. This integrated approach ensures maximum consistency and accuracy in the phenotypic data analysis.

### Spatial Transcriptomics (ST)

RNA was extracted from five-micron sections cut from the sample blocks chosen for the assay with the RNeasy Mini Kit (Qiagen). RNA quality was assessed by RNA integrity through measurement of DV200 value that had to be exceeding 50% measured using Agilent RNA 6OOO Nano Bioanalyseur (Agilent Technologies). In this study, we investigated the spatial distribution of gene transcripts within tissues using 10x Genomics’ Visium Spatial Gene Expression technology. Our methodology involved the following key steps: Sample preparation included obtaining two adjacent sections of five micrometers. The first slide underwent deparaffinization according to the Phenocycler protocol before Phenocycler treatment, while the serial slide designated for Visium only was exposed to a temperature of 60 degrees Celsius for two hours followed by deparaffinization. Finally, both sections were stained with H&E, followed by imaging and sequencing according to the Visium Spatial Gene Expression User Guide. Permeabilization, reverse transcription, second strand synthesis and cDNA amplification were performed using the Visium Spatial Gene Expression Reagent Kit (10X Genomics). Dual indexed libraries were made with the Library Construction Kit (10X Genomics) and Dual Index Kit TT Set A (10X Genomics) according to the manufacturer’s protocol. The final libraries were assessed using the Agilent Bioanalyzer High Sensitivity DNA kit and chip (Agilent Technologies). Loupe Browser 6.3.0 was used to estimate the capture area covered by the tissue within each frame on the slide to calculate the sequencing depths required. The libraries were pooled and sent for paired-end dual-indexed sequencing on the Nextseq 500 instrument (Illumina).

### Sequence alignment and annotation

Sequencing output and the histology images were processed using Space Ranger software v2.1.0 (10x Genomics). The Space Ranger mkfastq function was used for sample demultiplexing and to convert spatial barcodes and reads into FASTQ format. Space Ranger count function was used to align reads and calculate counts on the basis of the human reference genome (version GRCh38-3.0.0) and then align microscopic slide images and transcriptomes to generate barcode/UMI counts and feature spot matrices. Two sections of five µm were taken and multiplexed onto Visium Spatial gene expression slides (10x Genomics). Following incubation of the slides on an H&E staining library preparation, imaging and sequencing was performed in accordance with the Visium Spatial Gene Expression User Guide. FASTQ reads were mapped to the reference human genome (version GRCh38-3.0.0) and demultiplexed using SpaceRanger (v2.1.0). Using the 10X Loupe browser (v6.3.0). The raw sequenced read and expression data were processed and overlaid with the H&E image.

### ST data processing

Feature-barcode matrices for the *ST only* and *PP+ST* samples were imported into the R environment for quality control, normalization, dimensionality reduction and clustering. Firstly, data were loaded with the *STutility* package (now *semla*, Larsson *et al. Bioinformatics* 2023 PMID: 37846051) to filter out spots outside of tissue and for “manual” inspection of the images in order to remove spots over debris. Distribution plots for various technical metrics (feature counts, RNA counts) were exploratorily analyzed to observe and remove outliers. In addition, spots with under 200 genes detected and spots with >10% mitochondrial content were systematically removed. Filtered data were then normalized using the *SCTransform* function of the *Seurat* package (v4.3.0, Hao *et al. Cell* 2021 PMID: 34062119). UMAP plots of normalized expression data were generated for both samples based on the first 15 principal components (PCs), which were visually selected by inspecting Elbow plots of cumulative variance. Unsupervised graph-based clustering was performed on each dataset (ST only and PP+ST) separately using the *Seurat FindClusters* function. This approach implements a shared nearest neighbors (SNN) modularity based on a resolution value set to 0.4 for both datasets [best resolution: iterative tests of different resolutions for clustering and determine the most suitable one based on the stability of clustering produced]. Finally, samples *ST only* and *PP+ST* were merged and re-scaled for direct comparison.

### Malignant cell identification in ST data

We used SpaCET ^37^ to deconvolute spots in the Visium data. This algorithm estimates malignant cell fractions based on a gene pattern dictionary of copy number alterations (CNA) and malignant transcriptome signatures across tumors from the ∼10,000 patient samples spanning 30 tumor types from the Cancer Genome Atlas (TCGA). In each tumor ST data, SpaCET searches for malignant cell spots whose expression profiles correlate with the CNA or expression pattern of the relevant tumor type. SpaCET also allows to identify malignant cells in different spatial regions displaying distinctive expression profiles, which led us to label three malignant states (A, B, C). We then used Seurat’s FindMarkers function to identify which molecular features were associated to each of these malignant cell states.

### Pathologist annotations

High resolution H&E images of samples from the two slides of the biopsies of the same patient were provided to a pathologist for pathological and tissue annotation. The pathologist utilized the Loupe Browser to directly outline, and label various morphological features observed within the H&E tissue on the Visium spots format. Annotations were blindly performed (the pathologist was not made aware of analysis results) following all other analyses completed in this study.

### Spatial alignment of Phenocycler to Visium data

The alignment procedure involved identifying the optimal affine transformation using two parameters: 1) the scaling factor and 2) the shift term. The scaling factor was determined either directly from the microscopy images’ resolution metadata or estimated from the images themselves. The estimation protocol involved comparing the distribution of vertical and horizontal distances between the masked Visium and CODEX images. The shift term was determined by aligning the centers of mass of the two masked images.

These parameters were then fine-tuned using a grid search optimization procedure. The masked images were aligned, and the matching score was calculated using one of two alignment scores: 1) the total number of mismatched pixels in the aligned masks, or 2) the geometric mean of the number of mismatched pixels from each of the two masks independently.

### Generation of a pseudo-spot grid for Phenocycler data

The identified affine image transformation was applied to the cell centroids segmented from the Phenocycler data. The transformed centroid coordinates were then assigned to the coordinates of Visium spots by matching each centroid to the nearest spot within the pseudospot boundary (defined as a distance of less than 27.5 μm from the cell centroid to the spot center). The cell-level Phenocycler expression signals were then aggregated to the pseudo-spot level by summing the signals from all the corresponding cells. The identity of a pseudospot was defined by the most abundant cell type or state within each pseudospot. Finally, Phenocycler data at the pseudo-spot resolution was imported into the R environment and loaded as a *Seurat* object to allow direct comparisons with the Visium data.

## Data availability

This study’s raw sequencing data (fastq and BAM files) is under controlled access (patient data) and is available upon reasonable request to the corresponding author. The processed Visium gene expression matrices were stored in a Zenodo repository (10.5281/zenodo.13736222) that will be made public upon publication.

## Code availability

Software used for analysis is public and described in detail in the Methods section. Raw scripts and code are available at a GitHub repository linked to the article’s Zenodo page (10.5281/zenodo.13736222).

## SOX9 scoring

Adjacent tissue section to the slide used for Phenocycler and Visium analysis (PP+ST) was processed for immunohistochemistry to detect SOX9 expression. Immunohistochemical staining was performed on the Ventana Benchmark Ultra platform using an anti-SOX9 antibody [EPR 14335-78]. Chromogenic detection was achieved with DAB, resulting in a brown stain in SOX9-expressing cells. An H-score was calculated using the HALO software from Indica Labs. Briefly, after annotation of Malignant A and Malignant B/C areas, SOX9 intensity of expression was measured, and pixels were assigned a score between 0 and 3. The percentage of SOX9 positive tissue for each score was multiplied by the score value to calculate the H-score and compare malignant states A and B/C.

## CD70 Immunohistochemical Detection

A separate tissue section from the same tumor sample used for Phenocycler and Visium analysis (PP+ST) was processed for immunohistochemistry to detect CD70 expression. Immunohistochemical staining was performed on the Ventana Benchmark Ultra platform using an anti-CD70 antibody [E3Q1A]. Chromogenic detection was achieved with DAB, resulting in a brown stain in CD70-expressing cells.

