## Supplemental figures for "A new spatial multi-omics approach to deeply characterize human cancer tissue using a single tissue section"

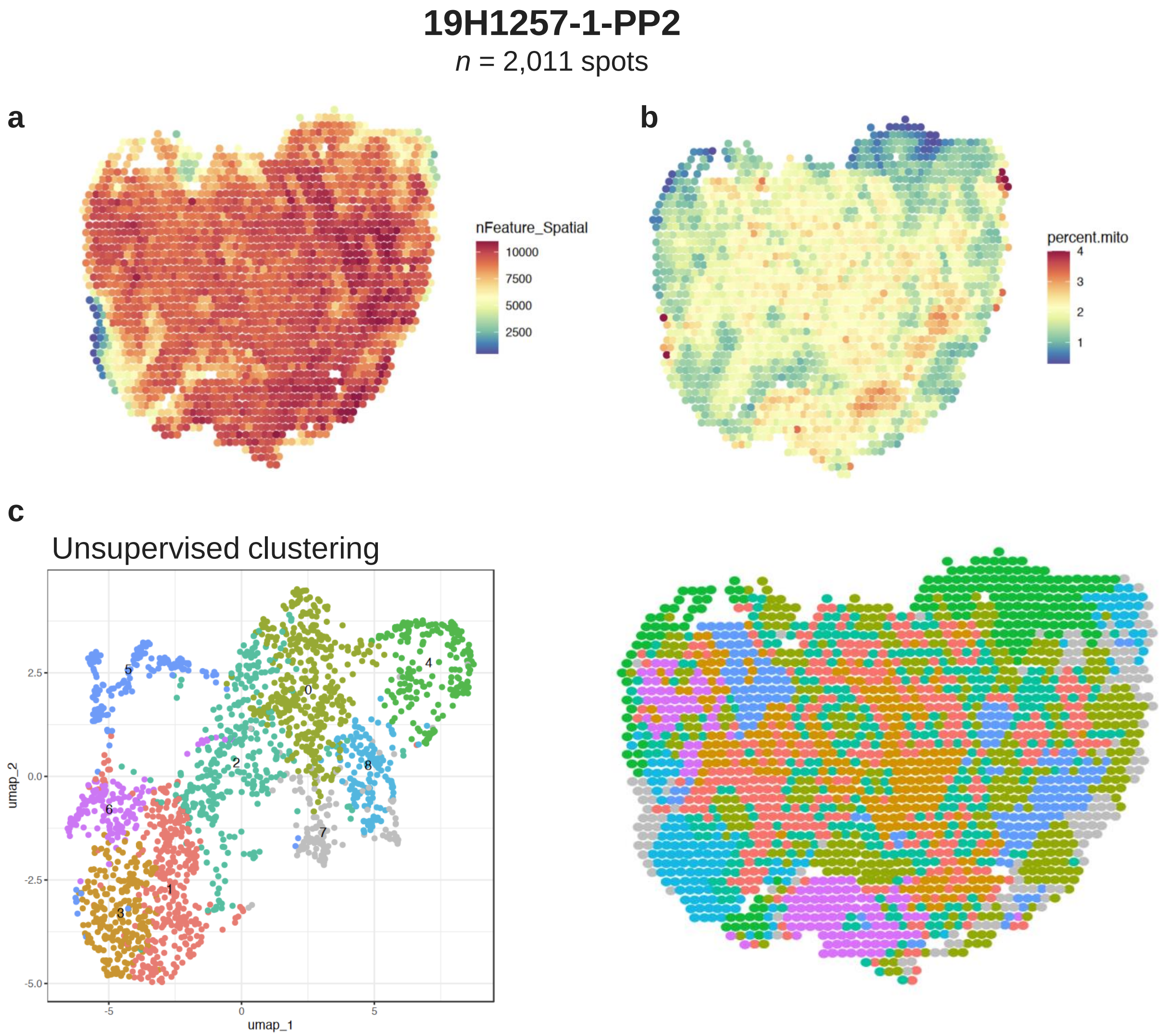


**Supp. Figure 1. Visium post-Phenocycler quality check on technical replicate** (a) Spatial distribution of gene features and UMI detection per spot in the technical replicate of 19H1257-1-PP2 (post-Phenocycler 2) sample. (b) Mitochondrial content comparison between the technical replicate and the original sample. (c) UMAP projection of transcriptomic data from 19H1257-1-PP2.

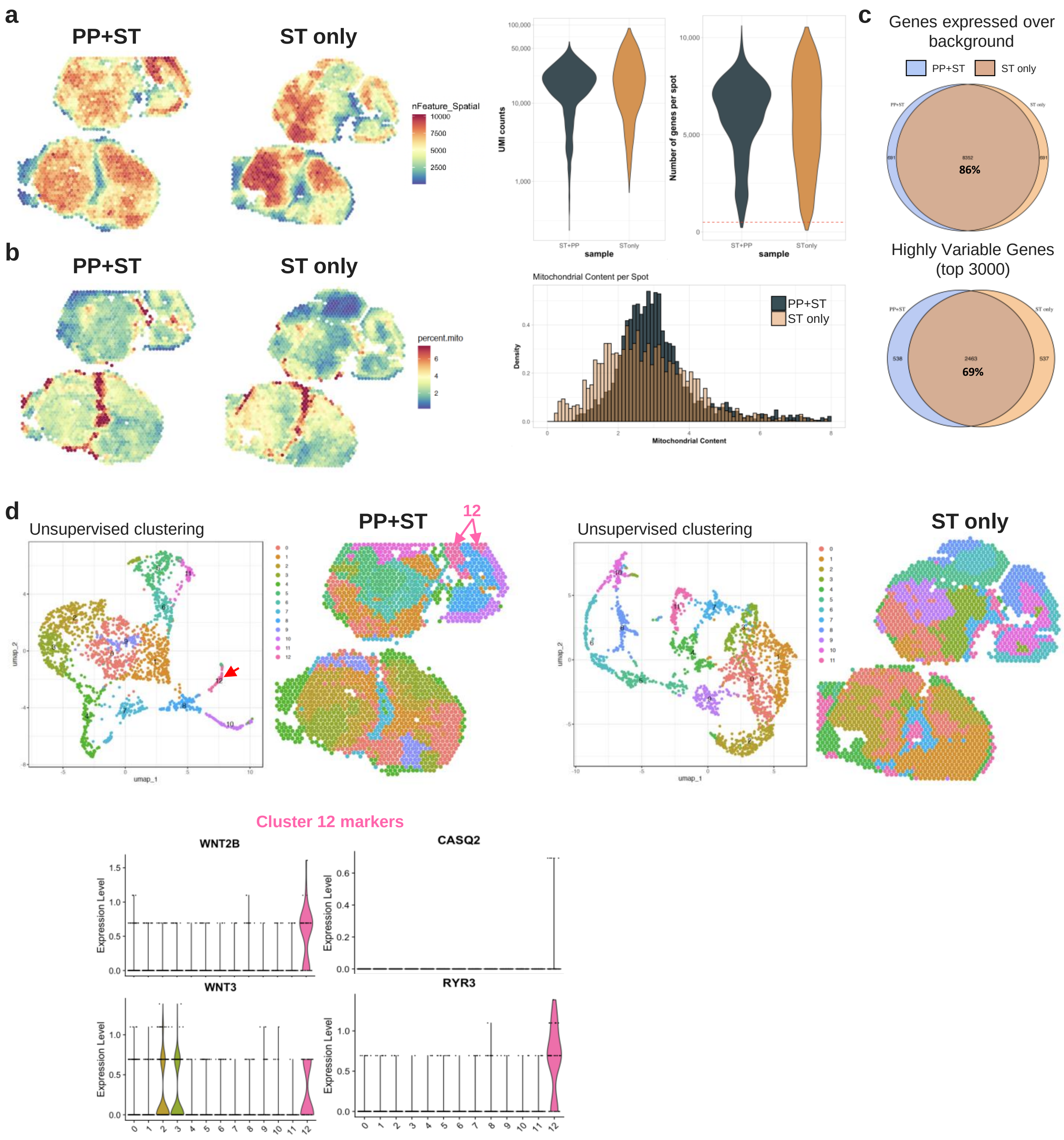


**Supp. Figure 2. Comparison of Visium post-PP vs Visium only on a new replicate.**
(a) Spatial distribution of gene features and UMI detection per spot in PP+ST and ST only conditions on another sample (17B5776-1). Violin plots represent the distribution of gene and UMI counts on a log10 scale for clarity. (b) Mitochondrial content per spot in PP+ST and ST only conditions. (c) Venn diagrams showing the overlap of genes expressed above background (top) and of the top 3000 highly variable genes (bottom) detected in PP+ST and ST only samples. (d) Transcriptomic landscape patterns derived from unsupervised clustering in PP+ST and ST only samples on the replicate. UMAP projection displays distinct clusters, with markers for cluster 12 shown at the bottom for specific genes (*WWTR1*, *CAV2*, *RYR3*, and *WNT7B*).

a

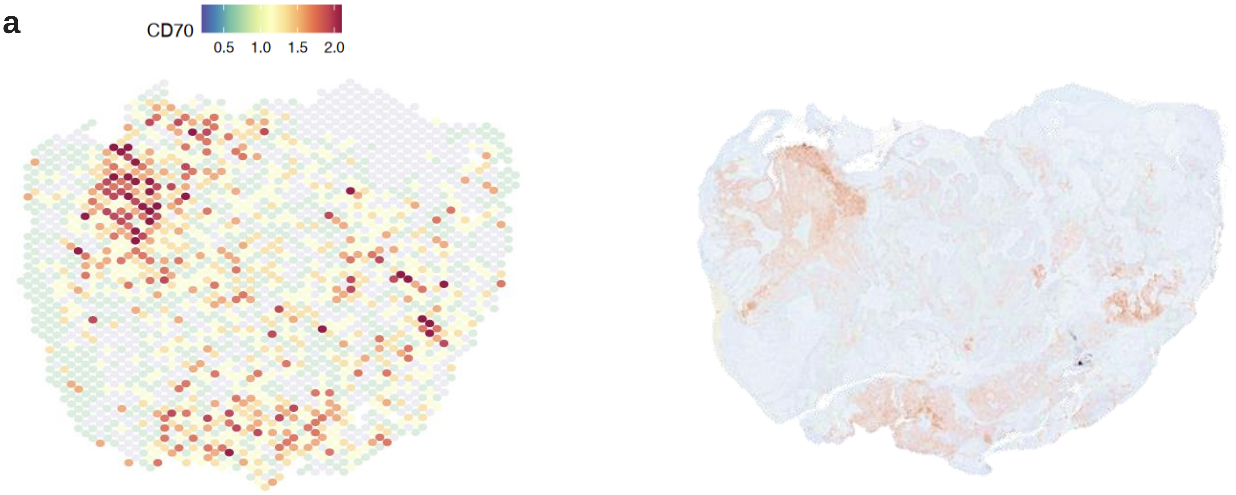


**Supp. Figure 3. Validation of CD70 expression by IHC** (a) Spatial distribution of *CD70* expression from Visium data and corresponding immunohistochemical staining using an antibody targeting CD70 protein on an adjacent slide.
